# Do combinations of clinical parameters related to caries activity status predict progression more accurately than individual parameters?

**DOI:** 10.1101/561878

**Authors:** I Floriano, R Matos, J Mattos-Silveira, ES Rocha, KR Ekstrand, FM Mendes, MM Braga

## Abstract

Few studies have addressed the predictive power of the clinical parameters used in assessing caries lesion activity. This study assessed the predictive validity of evaluating clinical parameters that are related to caries lesion activity status, individually and combined, in a long-term analysis. The occlusal surfaces of primary molars (1361 surfaces) were examined in 205 children according to the following clinical features: potential for plaque stagnation, colour, luster, cavitation, texture, and clinical depth. Cavities with frankly exposed dentine were excluded from this sample. After 1 year, 148 children (828 surfaces) were re-evaluated using the International Caries Detection and Assessment System to assess caries lesion progression. Progression was set as an outcome to verify the predictive power of the initially assessed clinical parameters. Different combinations of two or more parameters were also tested to check for any association with caries progression. Multilevel Poisson regression analyses were performed and the relative risk for each parameter/combination tested was calculated by considering a confidence interval of 95%. Forty percent of the reassessed surfaces presented caries progression after 1 year. Despite their surface integrity, dentine caries lesions were approximately 10-fold more likely and enamel lesions were approximately three-fold more likely to progress than sound surfaces. Similarly, cavitated lesions showed the highest risk of progression compared to sound/non-cavitated lesions. When only non-cavitated surfaces were considered, roughness proved to be a risk factor for caries progression. In conclusion, the lesions presenting clinical involvement of the dentine and even those cavitations clinically involving only the enamel had a higher risk of progression compared to sound or non-cavitated surfaces. For these lesions, the evaluation of other conjoint parameters seems unnecessary. Nevertheless, surface roughness can be a useful feature in predicting the risk of non-cavitated caries lesion progression.

## Introduction

Active enamel caries lesions have been generally defined as those that present rough enamel and loss of lustre (indicated by chalky-white enamel) [1, 2]. However, differential features linked to enamel caries activity status have been attributed to differences in enamel porosity and surface wear/polishing [3, 4]. These clinical features have traditionally been observed under specific conditions (areas of intense plaque accumulation and short-term remineralization of enamel lesions) and on specific surfaces [2]. Although the characteristics that guide caries lesion activity assessment are strongly intercorrelated [5], we hypothesized that some of the characteristics could be more closely related to caries progression than others, particularly considering the characteristics of occlusal surfaces.

Available visuo-tactile systems for caries lesion activity assessment recommend that the clinical features of caries lesions should be considered conjointly because caries is a dynamic process [6, 7]. Previous studies have verified the predictive validity of these systems [8, 9]. However, because these systems propose the conjoint evaluation of a pool of clinical characteristics detected through visuo-tactile inspection, to the best of our knowledge, we cannot affirm the predictive power of individual clinical characteristics associated with the status of caries lesions. Therefore, this is the first study to prospectively evaluate the influence of each clinical characteristic on caries lesion progression. Predictive validation has been indicated as the optimal choice for determining validity of the assessment of caries lesion activity [10], and determine how well a test (in this case, individual components of available systems) can predict further events [11], as caries progression.

Accordingly, we aimed to assess, on occlusal surfaces of primary molars, the predictive power of clinical features that have traditionally been used in caries lesion activity assessment. In addition, we also tested combinations of these variables to verify possible improvement in the prediction of caries progression.

## Materials and Methods

The Local Committee for Ethics in Research previously approved this research protocol (#01/06).

### Examiner Training

Two examiners were involved in this study: one responsible to the baseline examinations (MMB) and other, for the follow-up (IF). This last examiner (IF) was introduced to the ICDAS by an experienced examiner (MMB), engaged in previous clinical studies in caries diagnosis. This experienced examiner was considered as a reference examiner for this study. Initially, the original index description was studied, as previously reported [12]. The examiner then individually evaluated projections of clinical photographs of caries lesions. Finally, 36 occlusal surfaces of extracted primary molars were evaluated. After each phase of training, the reference examiner discussed any divergence between the examiners and also coordinated the training. The next step was initiated after all doubts and divergence had been solved.

### Participant Selection

Children were selected from those who had sought dental treatment at our dental school. All children who had at least one primary molar available to be examined were eligible for enrolment. The children’s oral assent and their parents’ written consent had to be obtained to guarantee participation in the study. Surfaces with restorations, hypoplastic defects, sealants, or frank cavities were excluded from the study sample. If a child presented more than one eligible molar, all of them could be included.

The required sample size was estimated based on the assessment of active caries lesions in children. Sample size calculation [13] was based on a prevalence of active lesions of 62.5%, observed in a previous study conducted on a Brazilian population [14], and a confidence level of 95%. For this calculation, we assumed one surface would be included per child. A minimum sample size of 126 surfaces was calculated and this number was increased by 20% to compensate for parent or children’s refusal to participate and for possible dropouts. Hence, a sample of 151 surfaces was required. Because more than one occlusal surface could be included in the sample, to compensate for the clustering effect, we assumed a factor of correction of 1.4. We determined that at least 212 surfaces were needed for our sample.

### Clinical Examination at Baseline

The children were examined in a dental unit and the examiners used a plane dental mirror, a ball-ended probe, and a three-in-one syringe. Before examination, teeth were gently cleaned with a rotating-bristle brush and pumice/water slurry.

In the baseline, an examiner experienced in caries diagnostic research (MMB) assessed eligible occlusal surfaces. For all assessments, a site was pre-selected for each surface by an external researcher on the basis of the highest ICDAS score found on the respective surface. Sites were recorded using a specific illustration in the participant’s file to guide next stages.

Sites were classified according to clinical characteristics of caries lesions that are generally understood to be associated with caries lesion activity status: potential of plaque stagnation on the basis of morphology and position of the surface, colour, lustre, surface integrity, depth, and texture [2]. The examiner did not use any specific system, but classified the sites as described in a previous report [5].

### Clinical Examination at the 1-year Follow-up

One year after the first examination, an examiner (IF), different from that responsible for baseline assessments, reassessed the children using the ICDAS [12]. In addition, restorations and teeth that had been extracted (because of caries) were also recorded. The clinical examination was performed under the aforementioned conditions. For this evaluation, the examiner followed pre-signaled charts with the occlusal sites evaluated at the baseline.

### Statistical Analyses

Caries progression was set as the outcome in further analysis and the evaluated sites were dichotomized into those that presented caries progression and those that did not. Each of the clinical features related to the activity status of caries lesions (potential for plaque stagnation, colour, lustre, surface integrity, texture, and lesion depth) was tested as an independent variable.

To evaluate caries progression, the baseline and follow-up assessment results were compared. Caries progression was considered when cavities with dentine exposure and/or teeth were restored or extracted because of caries as progression. Progression not related to cavitation exposing the dentine (e.g., ICDAS score 1– to ICDAS 2) was not considered for analyses. To verify the association of caries progression with the independent variables (clinical features), multilevel Poisson analyses were performed, considering the tooth and the child as the levels. Univariate analyses were performed both for the full sample and for non-cavitated lesions. For these analyses, we alternatively considered caries progression excluding cases that were restored after 1 year.

A multiple regression model was not used for the independent variables because of the likelihood of high collinearity among them. Indeed, one variable could cancel out the effect of another related variable in a multiple model if they are strongly associated with each other, despite being equally crucial individually in explaining the outcome. However, the interactions among some of the variables was tested to evaluate the possible benefit of combining these variables in assessing caries lesion activity.

Besides, to simulate the use of systems that combine these characteristics, we created other 2 independent variables regarding lesions activity status. First, we assumed that an initial or established active lesion would be whitish/yellowish, with no lustre and with rough enamel. If a lesion did not present these three features at the same time, it was classified as inactive. Second, we assumed that an active lesion should present at least two of the three aforementioned clinical features.

The relative risk for the clinical features (alone or combined with one or more other clinical features) was calculated, with 95% confidence interval (95% CI). The level of significance was set at 5%.

## Results

The intra-examiner reproducibility in the assessment of clinical parameters by the experienced examiner was high (Kappa values: 0.93 (95% CI = 0.90–0.97) for lesion depth and surface integrity, 0.98 (95% CI = 0.97–1.0) for plaque stagnation, 0.96 (95% CI = 0.93–0.99) for color, 0.88 (95% CI = 0.85–0.91) for texture, and 0.90 (95%CI = 0.87–0.93) for luster). Regarding the follow-up examiner, an intra-examiner agreement (weighted Kappa) of 0.849 and an inter-examiner agreement of 0.92, considering the experienced examiner, were reached.

Out of 205 children examined at the baseline (1361 surfaces), 100 were girls (49%) and 105 were boys (51%). The mean age (standard deviation, SD) was 7 (2.1) years. After approximately 1 year, 148 children (72%) were reassessed. The mean time of re-examination (SD) was 395 (70.8) days. In the follow-up, the sample comprised 70 girls (47%) and 78 boys (53%), with a total of 949 surfaces. From these, 828 occlusal surfaces that did not initially present frank cavitation into the dentine were reassessed. A total of 121 surfaces were unavailable for evaluation because the primary teeth were exfoliated.

The children reassessed after 1 year had similar caries experience on the basis of the decayed, missing, and filled surfaces index (mean dmfs + DMFS (SD) = 4.3 (0.9)) when compared with those who were not followed-up in the study (mean dmfs + 6 DMFS (SD) = 6.3 (6.7); *p* = 0.57). The number of reexamined caries-active children was also similar to those who were not followed-up (*p* = 0.90).

The status of examined surfaces at the baseline and at follow-up is shown in Table 1.

**Table 1.** Classification of the surfaces (n) at baseline, according to the clinical parameters, and at 1-year follow-up, according to the ICDAS [12].

| Clinical features |  | ICDAS (1-year follow-up) |  |  |  |  |  |  |  |  | Total |
| --- | --- | --- | --- | --- | --- | --- | --- | --- | --- | --- | --- |
|  |  | 0 | 1 | 2 | 3 | 4 | 5 | 6 | R | E |  |
| Potential for plaque stagnation | No | 5 | 1 | 1 | 0 | 0 | 0 | 4 | 5 | 0 | 828 |
|  | Yes | 314 | 166 | 51 | 113 | 21 | 45 | 23 | 65 | 14 |  |
| Colour | Absent | 262 | 55 | 6 | 10 | 2 | 4 | 3 | 4 | 1 | 828 |
|  | Dark | 8 | 33 | 11 | 25 | 10 | 12 | 5 | 10 | 5 |  |
|  | Whitish | 36 | 52 | 21 | 30 | 2 | 2 | 2 | 6 | 0 |  |
|  | Yellowish | 13 | 27 | 14 | 48 | 7 | 27 | 17 | 50 | 8 |  |
| Lustre | Present | 284 | 100 | 18 | 51 | 10 | 14 | 11 | 27 | 2 | 828 |
|  | Absent | 35 | 67 | 34 | 62 | 11 | 31 | 16 | 43 | 12 |  |
| Surface integrity | No | 3 | 12 | 6 | 40 | 14 | 34 | 20 | 55 | 13 | 828 |
|  | Yes | 316 | 155 | 46 | 73 | 7 | 11 | 7 | 15 | 1 |  |
| Texture | Smooth enamel | 288 | 112 | 20 | 51 | 6 | 11 | 5 | 20 | 1 | 828 |
|  | Rough enamel | 31 | 55 | 32 | 62 | 15 | 34 | 22 | 50 | 13 |  |
| Lesion depth | Sound | 260 | 56 | 6 | 10 | 1 | 5 | 3 | 4 | 1 | 828 |
|  | Enamel | 59 | 109 | 44 | 97 | 12 | 15 | 3 | 17 | 3 |  |
|  | Dentine* | 0 | 2 | 2 | 6 | 8 | 25 | 21 | 49 | 10 |  |

0 – 6 = ICDAS scores; E = indicated for extraction / extracted tooth due to caries; R = restored surface. \* lesions were clinically classified into dentine only if a shadow was observed under enamel (even without dentine exposure) – cavities exposing dentine were not considered in these analyses. The highlighted columns correspond to the surfaces on which were considered progression.

Dentine lesions had a probability of progression approximately 10 times higher than that of sound surfaces. Conversely, enamel lesions were only three times more prone to progression compared with sound sites (Table 2). Similarly, cavitated lesions were six times more closely associated with caries progression than non-cavitated surfaces. When only non-cavitated lesions were considered, dentine lesions (shadows) were strongly associated with lesion progression (Table 2).

**Table 2.**
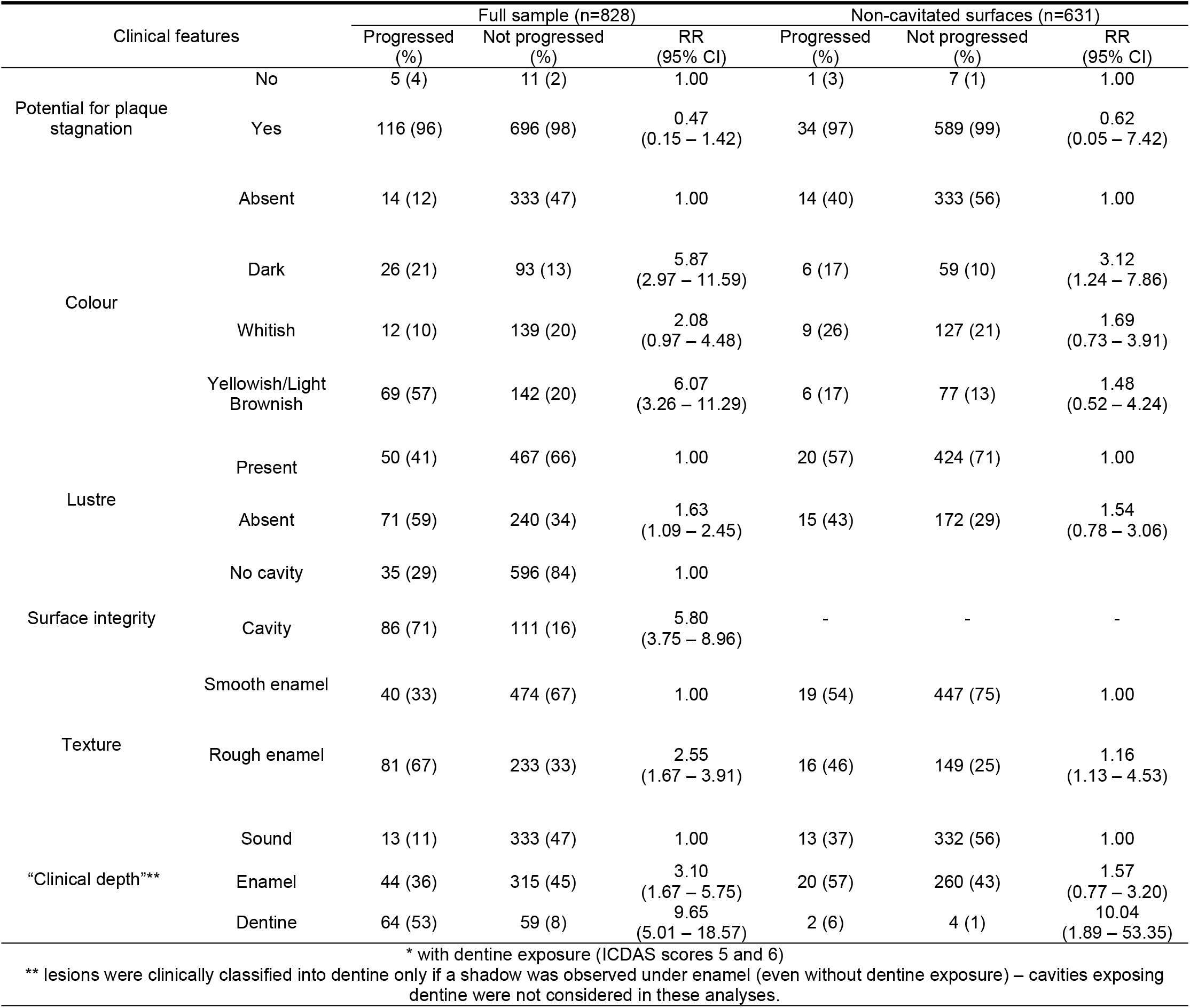
Relative risk (RR) with 95% confidence interval (95% CI) for caries progression (cavitation*, restoration or the tooth extraction due to caries) on occlusal sites examined followed by 1 year.

Colour is also a feature associated with caries lesion progression (Table 2). Brownish/black and yellowish sites were more likely to progress than sites without alteration in colour. Among the non-cavitated lesions, darker surfaces had a higher probability of progression compared to non-stained sites (Table 2).

Lesions without lustre progressed twice as high than those lesions with lustre. The same phenomenon was observed for surfaces presenting rough enamel compared to smooth enamel. However, for the non-cavitated lesions, only texture was a risk factor for caries progression (Table 2).

Similar trends were observed when restorations were not included in the outcome (Table 3). However, when only non-cavitated lesions were considered under these conditions, no associations were observed (Table 3).

**Table 3.**
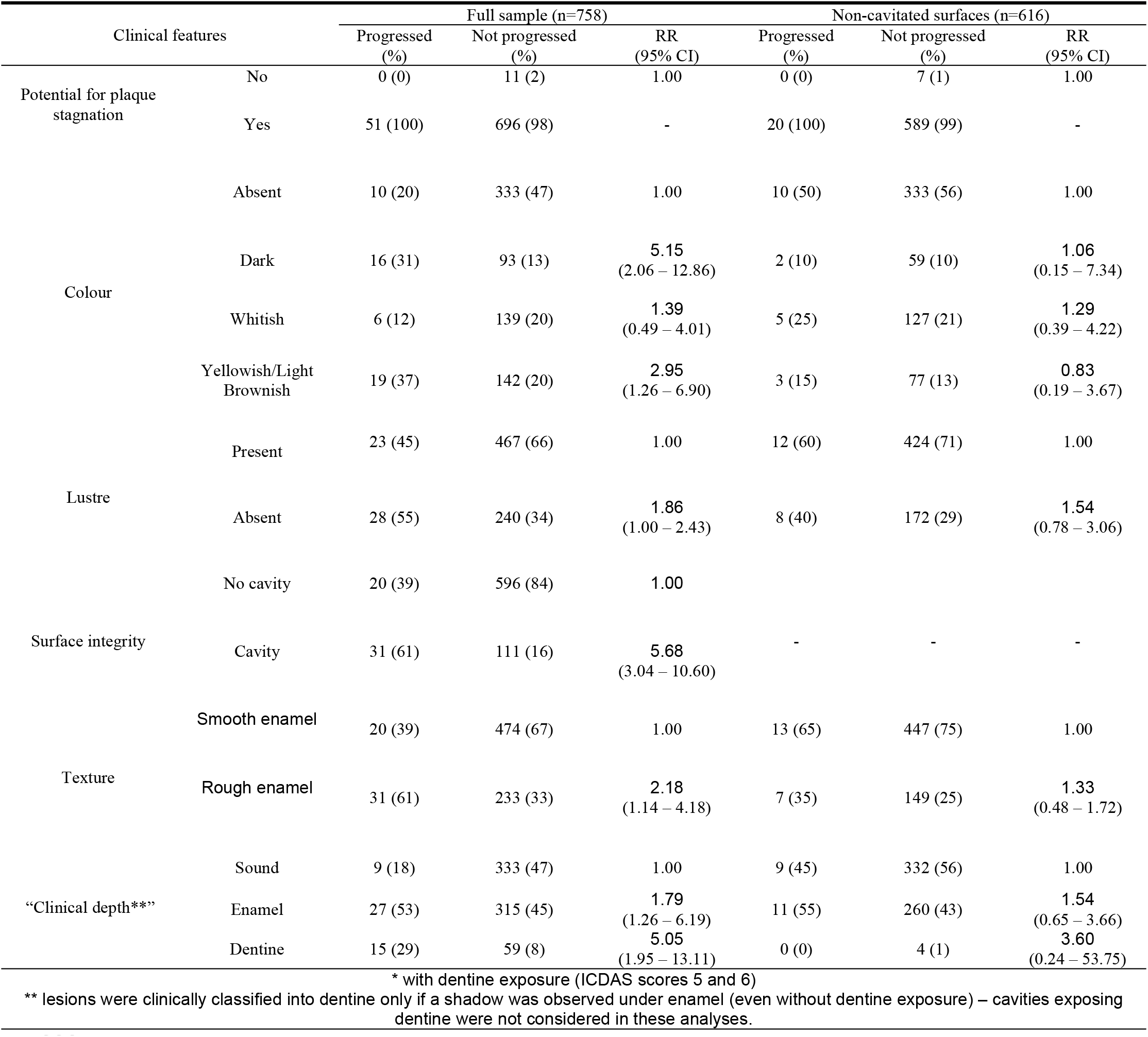
Relative risk (RR) with 95% confidence interval (95% CI) for caries progression (cavitation* or the tooth extraction) excluding possible restorations due to caries on occlusal sites examined followed by 1 year.

When combining colour, lustre and texture, similar risk of caries progression was observed, regardless of whether one or more parameters had been scored as positive for active status (Table 4). However, for non-cavitated lesions, when at least two clinical features were positive for active status, the sites had a two-fold higher risk for progression compared to sound surfaces (Table 4).

**Table 4.** Relative risk (RR) with 95% confidence interval (95% CI) for caries progression* after one year considering combination of clinical features related to caries lesion activity status on examined occlusal surfaces.

| Combination of clinical features |  | Full sample |  |  | Non-cavitated surfaces |  |  |
| --- | --- | --- | --- | --- | --- | --- | --- |
|  |  | Progressed (%) | Not progressed (%) | RR (95% CI) | Progressed (%) | Not progressed (%) | RR (95% CI) |
| **Combination 1: Texture and Colour** |  |  |  |  |  |  |  |
|  | No colour | 14 (12) | 333 (47) | 1.00 | 14 (39) | 333 (56) | 1.00 |
| Smooth enamel | Dark | 9 (7) | 56 (8) | 4.29<br>(1.85 – 9.92) | 3 (9) | 43 (7) | 1.90<br>(0.57 – 6.39) |
| Smooth enamel | Whitish | 3 (2) | 48 (7) | 1.56<br>(0.45 – 5.41) | 2 (6) | 44 (8) | 0.99<br>(0.22 – 4.45) |
| Smooth enamel | Yellowish | 15 (13) | 37 (5) | 5.78<br>(2.55 – 13.11) | 1 (3) | 27 (4) | 0.66<br>(0.07 – 6.23) |
| Rough enamel | Dark | 17 (14) | 37 (5) | 8.22<br>(3.75 – 18.05) | 3 (9) | 16 (3) | 5.66<br>(1.65 – 19.41) |
| Rough enamel | Whitish | 9 (7) | 91 (13) | 2.54<br>(1.10 – 5.88) | 7 (20) | 83 (14) | 2.12<br>(0.85 – 5.33) |
| Rough enamel | Yellowish | 54 (45) | 105 (15) | 6.81<br>(3.55 – 13.05) | 5 (14) | 50 (8) | 2.00<br>(0.64 – 6.20) |
| **Combination 2: Texture and Clinical Lesion “Depth”** |  |  |  |  |  |  |  |
|  | Sound | 13 (11) | 333 (47) | 1.00 |  |  |  |
| Smooth enamel | Enamel | 15 (12) | 134 (19) | 2.57<br>(1.22 – 5.40) |  |  |  |
| Smooth enamel | Dentine | 12 (10) | 9 (1) | 11.43<br>(4.60 – 28.37) |  | - |  |
| Rough enamel | Enamel | 29 (24) | 181 (26) | 3.51<br>(1.81 – 6.82) |  |  |  |
| Rough enamel | Dentine | 52 (43) | 50 (7) | 9.24<br>(4.66 – 18.32) |  |  |  |
| **Combination 3: Texture and Surface Integrity** |  |  |  |  |  |  |  |
| Smooth enamel | No cavity | 19 (16) | 447 (63) | 1.00 |  |  |  |
| Smooth enamel | Cavity | 21 (17) | 27 (4) | 9.49<br>(4.85 – 18.55) |  |  |  |
| Rough enamel | No cavity | 16 (13) | 149 (21) | 2.64<br>(1.37 – 5.08) |  |  |  |
| Rough enamel | Cavity | 65 (54) | 84 (12) | 8.24<br>(4.66 – 14.58) |  |  |  |
| **Combination 4: Texture and Lustre** |  |  |  |  |  |  |  |
| Smooth enamel | Lustre | 32 (26) | 423 (60) | 1.00 | 16 (46) | 402 (67) | 1.00 |
| Smooth enamel | No Lustre | 8 (7) | 51 (7) | 1.69<br>(0.75 – 3.81) | 3 (9) | 45 (8) | 1.30<br>(0.36 – 4.77) |
| Rough enamel | Lustre | 18 (15) | 44 (6) | 3.46<br>(1.80 – 6.65) | 4 (11) | 22 (4) | 3.15<br>(0.92 – 10.75) |
| Rough enamel | No lustre | 63 (52) | 189 (27) | 2.70<br>(1.68 – 4.32) | 12 (34) | 127 (21) | 2.12<br>(0.99 – 4.57) |
| **Combination 5: Surface Integrity and Lustre** |  |  |  |  |  |  |  |
| No cavity | Lustre | 20 (17) | 172 (24) | 1.0 |  |  |  |
| No cavity | No lustre | 15 (12) | 424 (60) | 1.75<br>(0.91 – 3.38) |  |  |  |
| Cavity | Lustre | 30 (25) | 68 (10) | 7.62<br>(4.14 – 14.02) |  | - |  |
| Cavity | No lustre | 56 (46) | 43 (6) | 6.90<br>(3.90 – 12.20) |  |  |  |
| **Combination 6: Texture, Lustre and Colour (I)** |  |  |  |  |  |  |  |
| No lesion |  | 13 (11) | 333 (47) | 1.00 | 13 (37) | 332 (56) | 1.00 |
| One or two clinical features positive for active status |  | 59 (49) | 173 (24) | 4.50<br>(2.47 – 8.19) | 13 (37) | 147 (25) | 1.71<br>(0.78 – 3.75) |
| Three clinical features positive for active status |  | 52 (43) | 170 (24) | 4.44<br>(2.38 – 8.28) | 9 (26) | 118 (20) | 1.95<br>(0.84 – 4.53) |
| **Combination 7: Texture, Lustre and Colour (II)** |  |  |  |  |  |  |  |
| No lesion |  | 13 (11) | 333 (47) | 1.00 | 13 (37) | 332 (56) | 1.00 |
| None or only one clinical feature positive for active status |  | 51 (42) | 466 (66) | 4.06<br>(2.06 – 7.99) | 20 (57) | 454 (76) | 1.10<br>(0.36 – 3.34) |
| At least two clinical features positive for active status |  | 81 (67) | 260 (37) | 4.65<br>(2.59 – 8.36) | 17 (48) | 182 (30) | 2.09<br>(1.01 – 4.33) |
| * it was considered as outcome the occurrence of cavitation, restoration or the tooth extraction. |  |  |  |  |  |  |  |
| I: To be active, the initial or established caries lesions should present all three characteristics |  |  |  |  |  |  |  |
| II: To be active, the initial or established caries lesions should present at least two of these three characteristics |  |  |  |  |  |  |  |

Cavitated lesions showed higher risk of progression regardless of texture or lustre (Table 4). Different from smooth surfaces, rough whitish lesions were more prone to progression than sound sites (Table 4). The texture evaluation of black/brownish caries lesions also seemed to improve the prediction of caries progression. Furthermore, the magnitude of the association with caries progression increased when texture and color were combined (Table 4). Among non-cavitated lesions, only rough black/brownish samples were associated with caries progression (Table 4). Additionally, no combination between luster and texture was associated with caries lesion progression.

## Discussion

Although available visuo-tactile systems for caries lesion activity assessment recommend evaluating the clinical features of caries lesions conjointly [7, 15], most of these characteristics have been individually associated with caries progression. When combinations of some of them were prospectively assessed, an additional contribution/benefit was observed only in specific situations.

This study aimed to clarify the predictive power of these clinical characteristics of active lesions for caries progression. Hence, we used a sample selected from children who had sought dental treatment. Because our sample was calculated a priori, we based it on a younger population [14]; older children may have less active caries lesions [16] because some lesions have more time and opportunity to be arrested and, accordingly, a larger sample would be necessary if we considered this difference. Conversely, the age group included may reflect a greater likelihood of seeking treatment, being representative of the population that we aimed to study. In addition, even considering this limitation, we obtained statistical power for demonstrating some crucial associations in our findings.

Similar trends were observed both when restorations were and were not considered as caries progression. Patients were followed-up but not treated by the researchers. Therefore, it is reasonable to consider restoration as progression even if the tooth had been restored by another professional before the established follow-up. We could not eliminate the possibility of interference from individual professionals’ choice for operative treatment [17]. Although punctual variation in estimates was observed, confidence intervals showed that the variables behaved similarly independent of the outcome considered. Greater differences were observed for non-cavitated lesions, but likely because of sample power, rather than changes in the association itself. Some sound surfaces and initial enamel lesions at baseline were restored during the study. However, we observed that a similar proportion of these lesions progressed to advanced lesions (ICDAS scores 5 and 6) after 1 year, reinforcing the aforementioned points.

Our results evidenced that severity may be a strong factor for predicting caries lesion progression. Between 60% and 85% of cavitated or dentine caries lesions progressed after 1 year. Lesion depth and the presence of cavities were two strongly interrelated features indicating caries lesion severity. Dentine lesions tend to become cavitated more easily because of their specific structure and composition [18]. Because of a high level of infection of the enamel–dentine junction, cavitated lesions are often histologically active [2, 19] and are consequently more difficult to arrest than non-cavitated lesions.

Cavitated lesions become inactive only when control of the biofilm through daily toothbrushing is possible [20], such as in considerably small and accessible cavities or extensive decay. The first studies concerning these lesions were conducted on large cavities. Thus, they were easily cleaned through toothbrushing and could be arrested [2] because the conditions were favorable. Besides difficulties in controlling the local biofilm, several cavitated caries lesions, clinically located at the enamel, can also present some dentine involvement [21]. Caries lesions scored as ICDAS 3 or 4 present higher progression rates than those with lower scores [22], likely explaining why the combination of lesion severity assessment and other parameters, such as luster and texture, did not provide additional benefit in the prediction of caries progression. Thus, established decay (shadows and/or cavitated lesions) could be the first sign of caries progression, indicating the need for specific measures to stop this process.

Differently from established caries lesions, for which severity may be sufficient for predicting progression, assessment of some other features of non-cavitated caries lesions could be useful. Non-cavitated lesions that presented at least two clinical features scored as positive for active lesions (whitish/yellowish color, loss of luster, or rough surface) [2] had a higher risk of progression than sites with no lesions. However, the same was not observed for lesions in which only one of these factors was identified.

Furthermore, the use of multiple combined parameters has likely been advocated since we are assessing a dynamic process and we can find mixed or intermediate forms of caries lesion status [2]. This conjoint assessment permits the classification of lesion status according to the most clinical features suggested. In addition, our findings suggest that among non-cavitated caries lesions, the risk of progression is similar if all or most clinical features are positive for active status. This observation corroborates the transitional process of caries arrest.

However, we found some differences in the association of the clinical features of non-cavitated lesions and their progression to cavities. Differently from roughness, lustre was not associated with caries progression and evaluating the luster of a rough non-cavitated lesion in occlusal surface of primary molars does not help predict its progression. Roughness and loss of luster are biologically related to caries lesion formation/progression because they reflect surface alterations resulting from acid attack and the increase in superficial porosity caused by demineralization, respectively. Studies that have demonstrated the loss of lustre as a classical characteristic associated with active caries have mainly evaluated areas of intense plaque accumulation, as areas around orthodontic appliances. In these studies, appliances were removed during the study to permit the tooth cleaning and fluoride application was intensified to stimulate quick remineralization of the surface, resulting in a gain in its lustre [4, 23].

Clinically, especially in occlusal surfaces, the reversion of lesion status could be slower and less evident than in aforementioned conditions. In addition, differences in enamel porosity may impede the differentiation of caries lesions and other enamel defects [24]. We should also consider that changes in the clinical appearance of caries lesions may have been due to professional cleaning prior to examination. We believe, however, that the effect of this procedure would have been low because we assessed occlusal surfaces. Accordingly, to predict caries lesion progression, it seems accurate and simpler to assess only the texture of non-cavitated occlusal caries lesions, instead of assessing both parameters (texture and luster) together.

Although whitish and yellowish lesions have been more associated with active caries, darker lesions are expected to be related to inactive caries [15]. However, in the present study they exhibited a similar risk of progression. Despite being a feature considered in various systems for assessing caries lesion activity status, colour should not be evaluated alone [6]. Although many inactive lesions are often dark brownish or black lesions, the contrary was not often true, reinforcing the need for conjoint evaluation of colour and other clinical features, such as roughness. Nevertheless, particular attention should be paid to the texture of lesions because of its subjective nature, which complicates standardization [25].

Because clinical features tend to reflect caries lesion activity at the moment of the clinical examination, a static time point, the importance of evaluating some features conjointly is paramount. However, our findings suggest that some of these parameters could be more helpful in this task than others, which could simplify lesion activity assessment.

In conclusion, caries with clinical involvement of the dentine as well as cavitated caries lesions (even if, with clinical involvement of only the enamel) had a higher risk of progression compared to sound or non-cavitated surfaces. Thus, evaluating other conjoint parameters seems unnecessary. However, superficial roughness can be a useful feature to help in predicting the risk of caries lesions.

## Acknowledgments

The authors are thankful to Prof. Dr. José Leopoldo Ferreira Antunes for checking the statistical analyses for pertinence and correctness. The authors would also like to thank the participants of the Post-Graduation in Pediatric Dentistry Seminar of Dental School, University of São Paulo for their critical comments and Editage for the English revision of this manuscript. São Paulo Research Foundation FAPESP (2011/16415-0 and 2013/27206-8) and the National Council for Technological and Scientific Development – CNPq (448013/2014-2/ MMB’s and FMM’s Research Productivity Scholarship) supported this study. The funders played no role in the study design, data collection and analysis, decision to publish, or preparation of the manuscript.

